# Mapping whole-tissue spatial multi-omics at single-cell resolution with STEP

**DOI:** 10.1101/2024.11.22.624797

**Authors:** Zheqi Hu, Zirui Zhu, Linghan Cai, Yangen Zhan, Xiangming Yan, Zhikang Wang, Jingyun Chen, Bingqian Sun, Songhan Jiang, Sijing Du, Hongpeng Wang, Xiangtao Li, Fang Wang, Guohua Wang, Qi Zou, Yixuan Du, Zhiyuan Yuan, Yongbing Zhang

**Affiliations:** School of Computer Science and Technology, Harbin Institute of Technology (Shenzhen), Shenzhen, China; Institute of Science and Technology for Brain-Inspired Intelligence, Center for Medical Research and Innovation, Shanghai Pudong Hospital, Fudan University Pudong Medical Center, Fudan University, Shanghai, China; Faculty of Computing, Harbin Institute of Technology, Harbin, China; School of Artificial Intelligence, Jilin University, Changchun, China; AI for Life Science Lab, Tencent, Shenzhen, China; Shanghai Academy of Artificial Intelligence for Science, Shanghai, China; Pengcheng Laboratory, Shenzhen, China

## Abstract

Spatial omics technologies have revolutionized our ability to map transcriptomic, proteomic, or metabolomic profiles within tissue contexts. However, existing spatial multi-omics approaches emphasize clustering or domain identification from measured profiles, while a general framework for whole-slice mapping in measured and unmeasured areas at single-cell resolution across additional omics layers remains lacking. Here we introduce STEP, a computational framework that integrates probabilistic inference with deep learning for flexible integration of histology, spatially resolved transcriptomics (SRT), single-cell, and/or multi-omics data. We validated STEP’s advantages across diverse applications. Furthermore, STEP enables tri-modal integration of Xenium, CITE-seq, and imaging mass cytometry to map transcriptome-wide gene expression and protein profiles from a small gene panel. Through quadra-modal integration of SRT, histology, single-nucleus RNA sequencing, and inCITE-seq, STEP reconstructs a 3D multi-omics brain cell atlas from six low-resolution SRT sections. Collectively, STEP provides a cost-effective tool for exploring spatial biology by integrating diverse, complementary modalities.

## Introduction

Spatial omics technologies have transformed our understanding of biological systems by mapping transcriptomic, proteomic, or metabolomic profiles in their native tissue context^1–3^. By preserving spatial information lost in non-spatial assays, they reveal cellular organization, function, and interactions. An ideal goal is to efficiently construct whole-tissue multi-omics maps across many sections at single-cell resolution. However, achieving this goal remains significantly challenging^4–6^.

Spatially resolved transcriptomics (SRT) has advanced fastest among spatial omics technologies, but current platforms remain limited^7–9^. Early sequencing-based SRT provides transcriptome-wide coverage but lacks single-cell resolution because each spot contains multiple cells^8^. Recent platforms achieve transcriptome-wide single-cell profiling^10,11^, yet some suffer from molecular lateral diffusion^12^ and others cannot cover entire tissue sections^13^. These shortcomings hinder the integration and joint analysis of such data with other spatial modalities, such as whole-slide spatially precise proteomics data. Notably, although some platforms claim subcellular resolution, analyses often aggregate signals at the cell level for robust interpretation (Supplementary Note 1). Imaging-based SRT offers single-cell resolution but requires predefined gene panels and lengthy cyclic imaging^8^. Single-cell RNA sequencing (scRNA-seq) provides comprehensive single-cell transcriptomes without spatial context^14^. Recent advances in spatial multi-omics technologies offer a promising route toward this ideal^15–18^, but achieving robust single-cell resolution remains challenging, and high costs and technical complexity limit widespread adoption. Computational integration of complementary single-modality data thus offers a practical, scalable way to approximate the ideal experiment^19^.

Existing computational approaches have primarily focused on the pairwise integration of these modalities. Gene imputation combines imaging-based SRT with scRNA-seq for transcriptome-wide coverage^20–22^, while deconvolution combines sequencing-based SRT with scRNA-seq to improve cell typing^23–25^. Other methods enhance SRT resolution to the subspot level^26^, predict gene expression from hematoxylin and eosin (H&E) images at spot or pixel resolution^27,28^, or infer cellular composition and expression from SRT data^29,30^. Many of these approaches do not fully exploit complementary information across modalities, limiting comprehensive multi-modal inference. Multi-omic spatial-domain identification methods^31,32^ improve niche discovery, but do not infer missing modalities or generate whole-slice single-cell maps. Recent tri-modal efforts have demonstrated single-cell spatial multi-omics^33^ but often omit histology, a valuable source of morphological context. OmiCLIP^34^ partially addresses this gap by aligning histology with SRT through CLIP-based contrastive learning^35^, but remains limited to patch-level bi-modal integration without single-cell resolution or multi-omics fusion.

We developed STEP, a hybrid computational framework that integrates probabilistic inference with deep learning. Building on methods that jointly consider histology, SRT, single-cell, and/or multi-omics data^36,37^, STEP provides a generalizable, platform-agnostic framework for diverse spatial multi-omics integration scenarios. By combining morphology with spatial omics information, STEP links SRT measurements to individual cells and supports both sequencing- and imaging-based SRT. For sequencing-based SRT, STEP extends single-cell mapping from the sequenced area to the whole tissue section; for imaging-based SRT, it expands gene coverage from a few hundred genes to the whole transcriptome. In a tri-modal integration of Xenium, CITE-seq, and imaging mass cytometry (IMC) data, it infers transcriptome-wide gene expression and protein profiles in human breast cancer at single-cell resolution. In quadra-modal integration of SRT, histology, single-nucleus RNA sequencing (snRNA-seq), and inCITE-seq, STEP reconstructs a high-resolution 3D multi-omics brain cell atlas using only six sections measured by low-resolution SRT, enabling cost-efficient single-cell 3D mapping. Overall, STEP provides a flexible and generalizable framework for reconstructing high-resolution spatial multi-omics information across diverse experimental platforms and integration scenarios.

## Results

### 1. Overview of the STEP workflow

STEP is a hyper-modal computational framework that supports flexible integration of diverse spatially resolved transcriptomics (SRT) data with single-cell reference data to generate whole-slice maps at single-cell resolution with expanded omics layers (Fig. 1). It accommodates sequencing- and imaging-based SRT, optional histological features, and transcriptomic or multi-omic references (Fig. 1a–c). Depending on the input settings, STEP generates whole-transcriptomic or multi-omic maps at single-cell resolution across the whole slice (Fig. 1d).

**Fig. 1.**
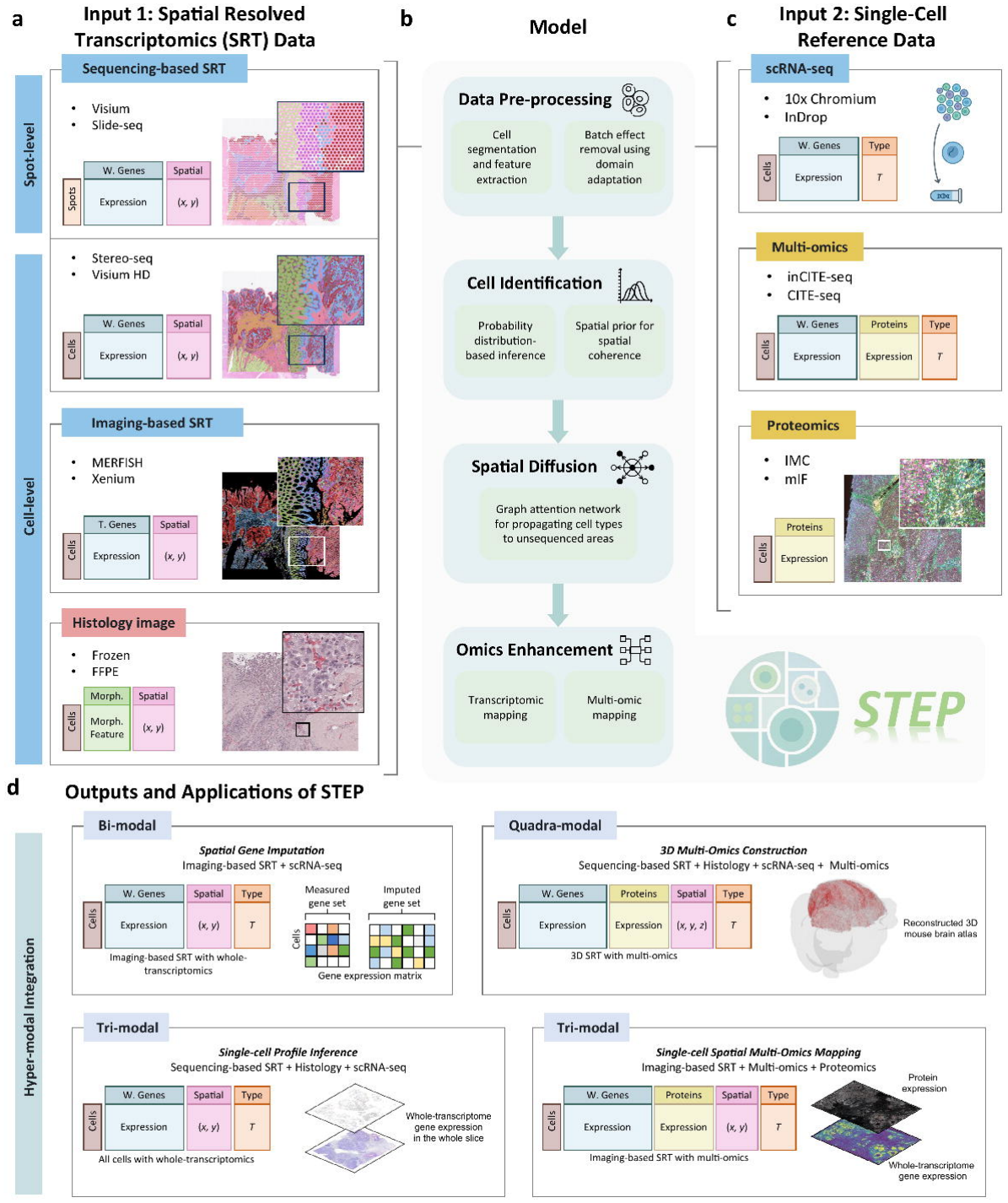
Overview of STEP. STEP is a hyper-modal computational framework that integrates diverse spatially resolved transcriptomics (SRT) data with single-cell references to refine whole-slice spatial profiles to single-cell resolution and expanded omics layers. **a** The first part of the STEP’s input data, including histological images, various spatially resolved transcriptomics data. **b** Data pre-processing and STEP’s main workflow, where the main workflow consists of three key stages: Cell Identification, Spatial Diffusion, and Omics Enhancement. **c** The second part of the STEP’s input data, including single-cell reference data. **d** Hyper-modal integrations enabled by STEP, including Spatial Gene Imputation for imaging-based SRT, Single-cell Profile Inference for sequencing-based SRT, Single-cell Spatial Multi-Omics Mapping, and 3D Multi-Omics Construction.

The STEP workflow comprises <u>Data Pre-processing</u> and three core modules: <u>Cell Identification</u>, <u>Spatial Diffusion</u>, and <u>Omics Enhancement</u> (Fig. 1b, Supplementary Notes 2–9 and Methods). <u>Data Pre-processing</u> performs specific operations on different data inputs, including cell segmentation^38^, optional morphological feature extraction from histology, informative gene selection^39^, and batch/platform effect removal using an adjusted conditional variational autoencoder (CVAE)^40^ (Supplementary Figs. 1 and 2). <u>Cell Identification</u> uses probabilistic inference to assign cell types to segmented cells within sequenced areas by linking cell types, gene expression, and optional morphological features (Supplementary Fig. 3). <u>Spatial Diffusion</u> uses a graph attention network (GAT)^41^ to propagate information to unsequenced areas, inferring cell types and informative gene expression across the whole slice (Supplementary Fig. 4). <u>Omics Enhancement</u> then uses scRNA-seq or single-cell multi-omics references to infer whole-transcriptome or multi-omic profiles, respectively (Supplementary Fig. 5).

STEP supports multi-modal integration by dynamically adjusting its module usage to accommodate specific modality combinations, including bi-modal, tri-modal and quadra-modal configurations (Fig. 1d and Supplementary Figs. 6–9). For sequencing-based SRT data, STEP infers cell-type annotations and recovers gene expression profiles in unsequenced areas, while for imaging-based SRT data, STEP does not invoke the <u>Spatial Diffusion</u> module, but instead directly performs <u>Cell Identification</u> and <u>Omics Enhancement</u>, yielding whole-transcriptomic maps at single-cell resolution. When single-cell multi-omic reference data are available, STEP executes multi-omic mapping following transcriptomic mapping during omics enhancement, delivering tissue-wide, single-cell-resolution multi-omic maps or even reconstructing 3D multi-omic atlases (Supplementary Notes 10–13 and Supplementary Table 1).

### 2. Histological cellular features guide single-cell mapping

Human Pancreatic Ductal Adenocarcinoma (PDAC) is widely studied to understand cellular heterogeneity and the tumor microenvironment. We collected data from two PDAC tissue slices^24,42^, PDAC-A and PDAC-B, generated by Spatial Transcriptomics (spot-level resolution, whole-transcriptome coverage, and matched H&E images), together with InDrop scRNA-seq profiles as the single-cell reference, to evaluate STEP’s performance in cell-type identification and gene expression prediction at single-cell resolution (Fig. 2a, Supplementary Fig. 10, Supplementary Note 14 and Methods).

**Fig. 2.**
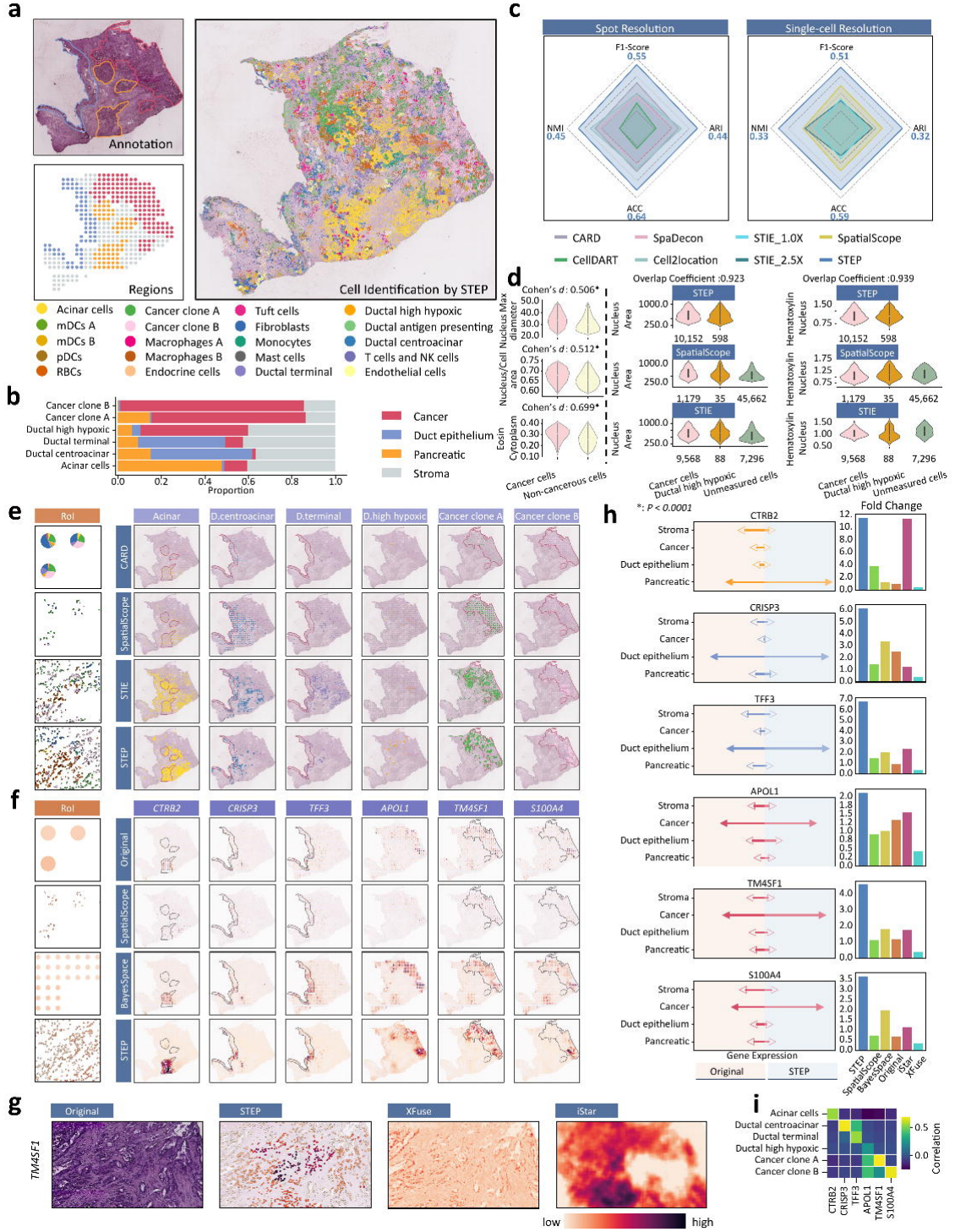
Cell identification and gene expression prediction at single-cell resolution by STEP. **a** Hematoxylin and eosin (H&E)-stained histological image of a pancreatic ductal adenocarcinoma section (PDAC-A) from Spatial Transcriptomics with manual pixel-level region labeling (top left) and spot-level labeling (bottom left). STEP’s cell identification results on PDAC-A sections (right). The legend for PDAC-A cell types is shared within Fig. 2. See panel **b** for the legend of region annotations. The remaining regions labeled at the pixel level were considered as Stroma. **b** Proportion of selected cell types in annotated regions inferred by STEP. **c** Radar plot comparing the performance of various models at the spot level and the single-cell level. STIE_1.0X and STIE_2.5X refer to the results of the STIE model when the spot diameter was enlarged to 1x and 2.5x, respectively. **d** Violin plots showing heterogeneous morphological features statistically classified by STEP, effect sizes (Cohen’s *d*) and corresponding *p*-values obtained by Mann-Whitney *U* Test (left). Violin plots through various methods and the number in the bottom right corner representing the number of cells in the violin plot (right). In STEP, the overlap coefficient shown above each panel quantifies the similarity of the morphological feature distributions between cancer cells and ductal high-hypoxic cells. “Unmeasured” refers to cells not covered by the algorithm, potentially containing the specified cell type. Center line represents median, lower and upper hinges represent first and third quartiles, and whiskers extend from hinge to ±1.5× interquartile range (IQR). **e** Visualization of cell identification results at different spatial resolution levels using representative methods and the same Region of Interest (RoI). **f** Visualization of various marker genes at different spatial resolution levels using representative methods and the same RoI displaying the unique molecular identifier (UMI) counts obtained by those methods. **g** Comparison of gene expression prediction by STEP and pixel-level regression methods within the same RoI. **h** Lollipop plots comparing the average expression of marker genes predicted by STEP versus the original gene expression in each region. The solid arrow indicates the highest enrichment value (left). Bar plots show the fold change of each marker gene in the expected domain versus other domains, where higher values indicate stronger domain specificity (right). **i** Heatmap showing the correlation between selected cell types and their corresponding marker genes at single-cell resolution, as inferred by STEP.

Since accurate cell-type identification at the single-cell level is crucial for analyses of SRT data, we benchmarked methods designed for similar tasks on the PDAC data, which fall into two categories: spot-level cell type deconvolution methods including SpaDecon^43^, CellDART^23^, CARD^24^, and Cell2location^25^, which predict the cell type proportions within each spatial spot, and single-cell-level cell type inference methods including STEP, STIE^30^, and SpatialScope^29^, which infer the cell type of each segmented cell (Methods). Following the benchmarking procedure described in Methods, we evaluated cell identification using tissue-region annotations and the known regional enrichment of cell types, with separate comparisons against spot-level deconvolution and single-cell-level inference methods. STEP achieved higher performance across multiple evaluation metrics in both comparisons (Fig. 2b,c, Supplementary Figs. 10–14 and Supplementary Notes 15 and 16).

We further qualitatively visualized cell identification results from different methods (Fig. 2e and Supplementary Figs. 15– 18). As shown in Fig. 2e and Supplementary Fig. 17, STEP provided accurate, fine-grained spatial distribution predictions of normal pancreatic regions with well-defined boundaries for acinar cells, which are responsible for the pancreas’s exocrine function. This advantage is attributed to STEP’s ability to model and leverage the gaps between sequenced spots, which preserves tissue continuity (Methods). Conversely, SpatialScope is confined to each spot, and STIE extends tissue coverage only to a limited degree (Fig. 2e, Supplementary Figs. 17 and 19). Additionally, ductal high-hypoxic cells are typically sparsely distributed within hypoxic microenvironments^44^. Consistently, in our results they accounted for only 1.21% of cells, making them challenging to detect using deconvolution methods (Fig. 2e, Supplementary Fig. 15). By contrast, the enhanced spatial resolution of STEP allows for precise characterization of these rare cells, successfully capturing their distribution along the periphery of cancerous areas (Fig. 2e). Note that STEP divided the cancer region into two sub-regions: an upper subregion dominated by cancer clone A cells and a lower subregion dominated by cancer clone B cells (Fig. 2e). Moreover, STEP demonstrated robustness in cell identification and showed resilience to experimental artifacts (e.g., staining artifacts), whereas STIE could not achieve (Supplementary Fig. 18). These findings collectively establish STEP as a reliable framework for cell identification, capable of capturing intricate spatial patterns and overcoming limitations inherent in other methods.

To evaluate whether cell identification by STEP is consistent with known morphological differences, we examined the distributions of morphological features across predicted cell types (Fig. 2d and Supplementary Fig. 20). We found that the morphological features of cancer cells identified by STEP were significantly different from those of other cell types, which aligns with the biological expectations (Fig. 2d and Supplementary Fig. 20). For instance, cancer cells are known to exhibit marked nuclear structure and chromatin distribution due to their altered proliferative and metabolic states^45,46^. These changes often result in nuclear enlargement and an increase in the maximum nuclear diameter, further manifestedby an increase in eosinophilic optical density in the cytoplasm (Cohen’s *d* = 0.699, *p* < 0.0001) and a higher nucleus-to-cell These changes often result in nuclear enlargement and an increase in the maximum nuclear diameter, further manifested area ratio (Cohen’s *d* = 0.512, *p* < 0.0001) (Fig. 2d and Supplementary Fig. 20). Additionally, the morphological feature distribution of rare ductal high-hypoxic cells, as determined by STEP, closely mirrors that of cancer cells with a similar trend in the Nucleus Area (92.3% overlap coefficient) and Hematoxylin optical density in the Nucleus (93.9% overlap coefficient), which is consistent with the prevailing biological consensus^47–49^ (Fig. 2d).

Since STEP not only identifies cell types at the single-cell level but also predicts gene expression at single-cell resolution, we further benchmarked its performance in super-resolution gene expression prediction against BayesSpace, SpatialScope, iStar, and XFuse (Fig. 2f and Supplementary Figs. 21 and 22). Among these methods, BayesSpace^26^ primarily subdivides spots into multiple sub-spots, while SpatialScope is limited to predicting gene expression of individual cells within a given spot (Supplementary Fig. 22). iStar’s^27^ results lack cell-level features and suffer from significant blurring artifacts, while XFuse^28^ can identify cell outlines but has low accuracy in gene prediction (Fig. 2g). This blurring presumably stems from their image-to-gene regression frameworks that average signals across entire patches without explicit cell-instance constraints (Fig. 2g). STEP addresses these limitations by providing fine-grained predictions of gene expression at the single-cell level across the whole slice (Fig. 2f). This characteristic allows for more comprehensive spatial mapping of biomarkers, facilitating the exploration of complex cellular microenvironments.

We further assessed the spatial enrichment of representative marker genes and their correspondence with predicted cell types, supporting the spatial and cell-type consistency of STEP’s gene expression predictions (Fig. 2h,i and Supplementary Note 17).

Beyond PDAC, we additionally constructed Visium-like pseudo-spots from single-cell-resolution Xenium datasets of human breast cancer and human colorectal cancer. These supplementary benchmarks enabled direct comparison against the original Xenium profiles and annotations, and showed that STEP maintained improvement in both cell identification and gene prediction across additional cancer contexts (Supplementary Figs. 23–29).

### 3. Robust cell identification across diverse settings

Human breast cancer is a prevalent disease with significant clinical impact, making it a critical focus for research across multiple platforms and modalities^50^. Therefore, we selected the BRCA-A dataset generated by 10x Visium (spot-level resolution, whole-transcriptome coverage, and matched H&E images), together with 10x Chromium scRNA-seq profiles^51^ as the single-cell reference, to validate STEP’s robustness in cell identification.

First, we evaluated STEP’s robustness to different scRNA-seq references. Building upon the 10x Chromium reference used above, we further incorporated two additional breast cancer scRNA-seq references: an independent 10x Chromium reference^52^ and one reference generated using the InDrop platform^53^ (Fig. 3a–c, Supplementary Figs. 30–34, Supplementary Note 18 and Methods). Despite differences in cell-type composition and gene expression patterns across these three single-cell references, STEP still clearly distinguishes between the various cell types (Fig. 3a–c and Supplementary Fig. 35). Using the InDrop single-cell reference, STEP predicted an enrichment of Prolactin-Inducible Protein (PIP)-expressing mammary luminal cells in cancerous regions, with 92.3% of these cells located within cancer regions (Fig. 3c). Notably, PIP is a clinically relevant breast cancer marker associated with well-differentiated, less aggressive tumors, underscoring translational relevance^54^. Consistently, analyses based on the independent 10x Chromium single-cell reference (Methods) assigned cancer-associated populations to tumor regions and immune-associated populations to immune regions (Supplementary Fig. 35).

**Fig. 3.**
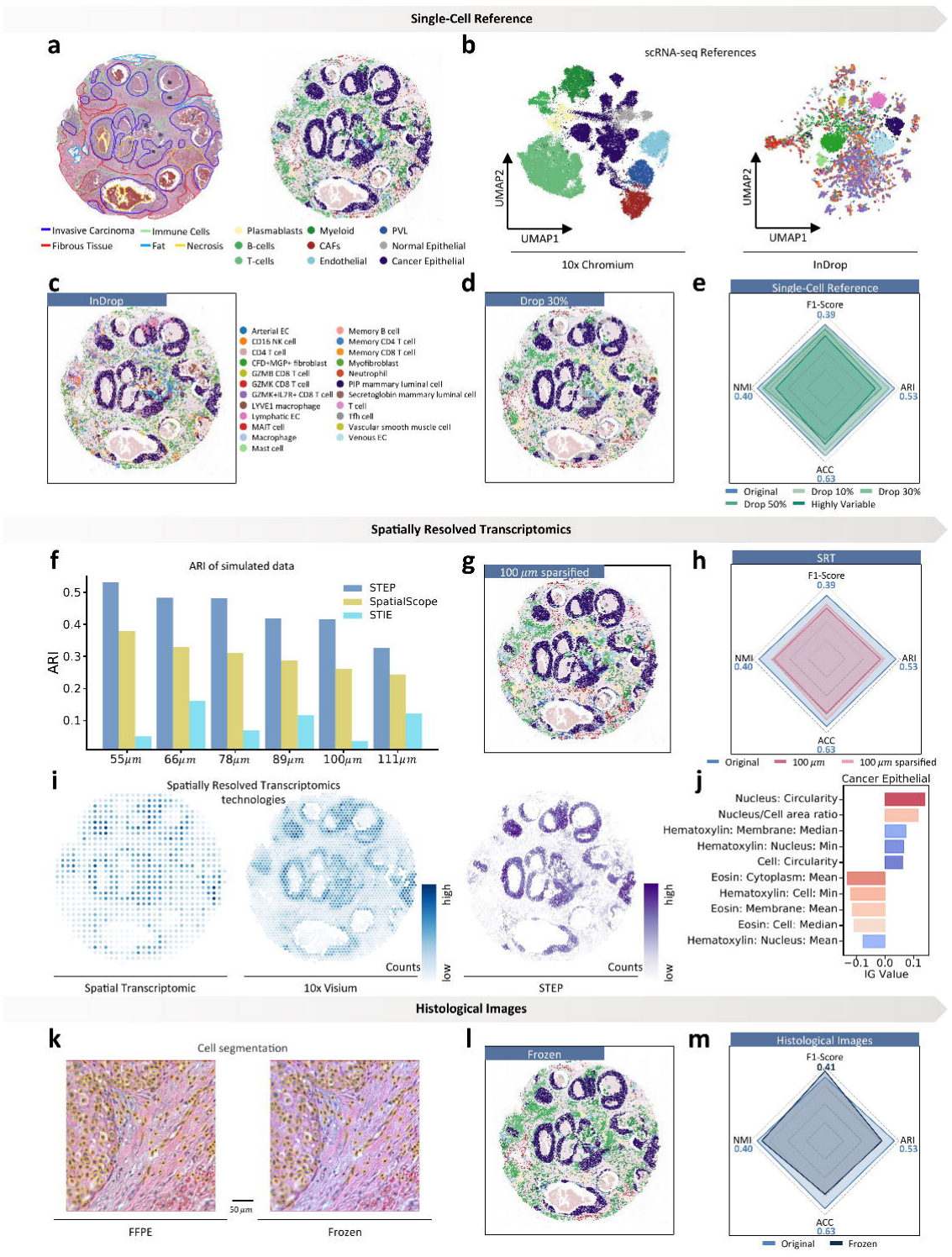
Robustness evaluation for STEP on different scenarios. **a Hematoxylin and eosin (H&E)**-stained histological image of human breast cancer section BRCA-A from the 10x Visium platform with manual pixel-level region labels (left) and STEP’s cell identification results (right). Cancer cells are expected to be enriched in cancer regions, while T cells, B cells, and myeloid cells are expected to be localized in immune regions. The legend for BRCA-A cell types is shared within Fig. 3. **b** UMAP visualization of single-cell RNA sequencing (scRNA-seq) data from breast cancer tissue using the 10x Chromium and inDrop platforms. **c** STEP’s cell identification results using scRNA-seq data collected with the inDrop platform. The legend is shared with **b. d** STEP’s cell identification results after randomly dropping 30% of the selected genes. **e** Radar plot comparing the performance of informative gene selection methods on scRNA-seq data. **f** Bar plot of Adjusted Rand Index (ARI) performance on simulated spatially resolved transcriptomics (SRT) data of different resolutions. **g** Cell identification results on simulated Spatial Transcriptomic platform’s data. **h** Radar plot comparing the performance of different spatially resolved transcriptomics data. **i** Scatter plot displaying the unique molecular identifier (UMI) counts at each spatial location for both the simulated Spatial Transcriptomic and 10x Visium platforms (left). The UMI counts at single-cell resolution predicted by STEP (right). The color scale is normalized to the 0-1 range. **j** Morphological features of cancer epithelial cells used as contributing factors for cell identification. The Integrated Gradients algorithm was employed to extract the contribution of morphological features to cell recognition, with the top contributing features for cancer epithelial cells selected. **k** Comparison of cell segmentation between the virtual frozen and original sections within the same Region of Interest (RoI). **l** STEP’s cell identification results on the virtual frozen section. **m** Radar plot comparing the performance between different histological image styles.

Next, we evaluated STEP’s robustness to interventions of the single-cell reference. We first performed a comparative analysis against random gene-drop on a single-cell reference (Supplementary Fig. 36). STEP retained over 92.0% of its original performance even after removing 30.0% of the genes, confirming that our gene selection strategy based on differentially expressed genes (DEGs)^39^ enables effective utilization of global expression heterogeneity (Fig. 3d,e and Supplementary Table 2a). Building on this, we conducted additional robustness analyses on the single-cell reference by merging, removing, subdividing, or unbalancing selected cell types (Methods). Across these interventions, STEP maintained consistent delineation of functional regions (Supplementary Figs. 37 and 38), demonstrating robustness to reference composition. Collectively, these results validate STEP’s compatibility with both 10x Chromium and InDrop references, support reliable cross-platform predictions at the cell-type and regional levels, and highlight its potential to facilitate spatially resolved studies in breast cancer.

As STEP also takes SRT data and histological images as inputs, we evaluated its robustness to variation in both modalities. Using simulated BRCA-A data with varying spot sizes and sparsity, we found that STEP maintained robust cell identification across SRT resolutions (Fig. 3f–i and Supplementary Note 19). STEP also maintained robust cell identification across histological styles and feature representations in the BRCA-A dataset (Fig. 3j–m and Supplementary Note 20).

### 4. Revealing tumor heterogeneity in unsequenced tissue

While tumor tissues can be analyzed with SRT, the field of view covered by current platforms is often limited, leaving substantial regions with only histological images. Inferring gene expression in these unsequenced areas from sequenced areas is therefore important, as it enables more accurate modeling of tumor heterogeneity and microenvironmental interactions, which are critical for understanding cancer progression and therapeutic resistance.

Using spot masking and transfer experiments on human breast cancer sections, we evaluated STEP’s ability to reconstruct spatial profiles in unsequenced tissue. STEP maintained cell identification and gene expression prediction under reduced spatial sampling and enabled reconstruction across adjacent sections and different patients (Fig. 4a–g, Supplementary Figs. 48–60 and Supplementary Notes 21 and 22).

**Fig. 4.**
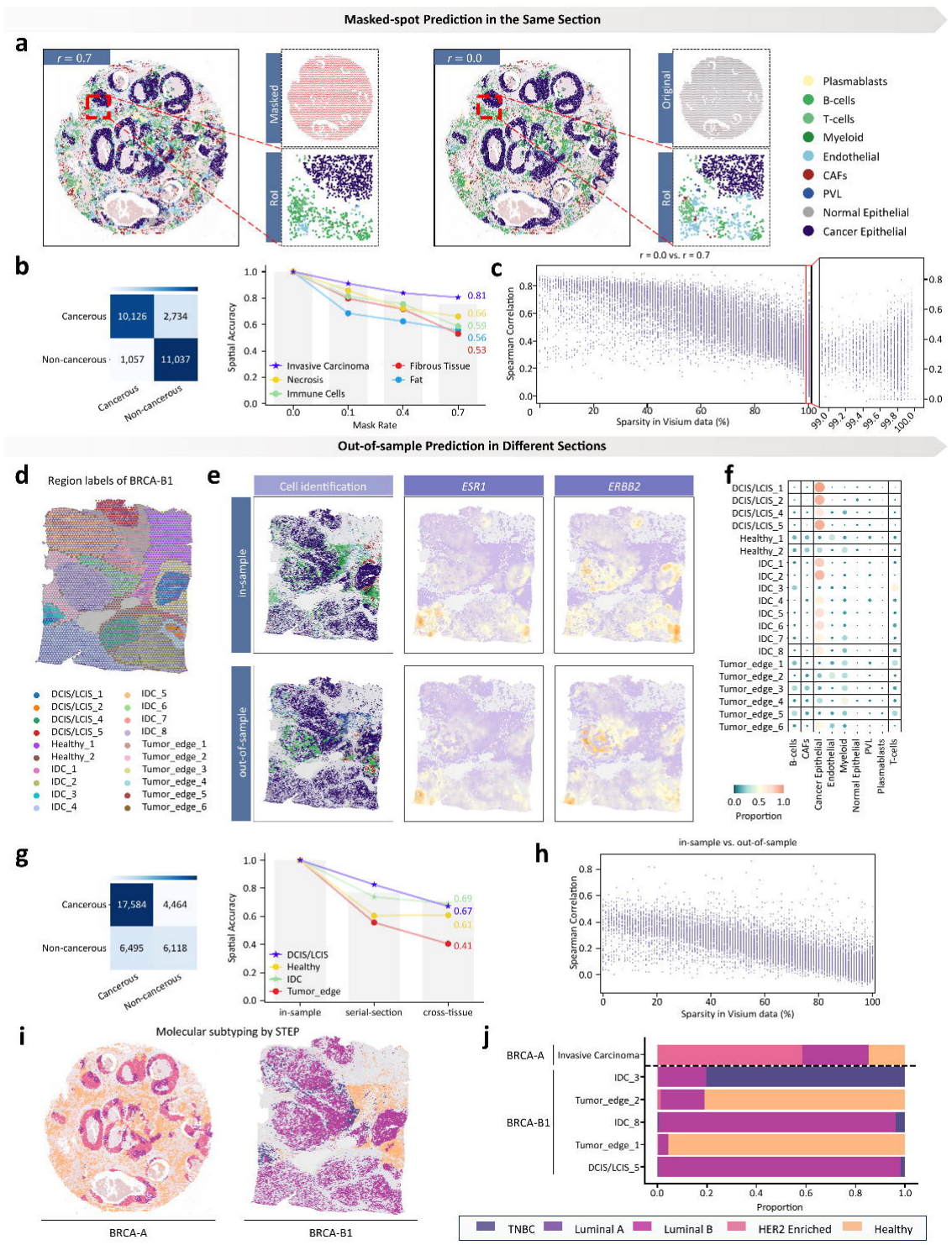
Reconstruction of spatial gene expression in unsequenced tissue with STEP. **a** Experimental results of STEP with mask rate *r* = 0.7(left) and the original data *r* = 0.0(right). In the same Region of Interest (RoI), the model shows superior accuracy in identifying Cancer Epithelial cells, while the accuracy of subtype classification in normal cells is relatively lower. In the upper right diagram, masked spots are shown in red, while retained spots are shown in gray. **b** Confusion matrix comparing the results of the masked data with the original data. Non-cancerous cells: Plasmablasts, B-cells, T-cells, Myeloid cells, Endothelial cells, cancer-associated fibroblasts (CAFs), perivascular-like (PVL) cells, and Normal Epithelial cells. Cancerous cells: Cancer Epithelial cells (left). The spatial accuracy (sACC) per region under different mask rates is shown in a line graph, with stars indicating cancerous regions and circles indicating non-cancerous regions. The sACC computed over all regions under different mask rates is displayed in a bar plot (right). **c** Spearman correlation between the single-cell gene expression results using the masked data and the original data for genes, along with their corresponding measurement sparsity. **d** Hematoxylin and eosin (H&E)-stained histological image of human breast cancer section BRCA-B1 from the 10x Visium platform with manual region labels. **e** Visualization of cell identification and marker gene prediction on BRCA-B1 section, along with serial-section transferred results from BRCA-B2. **f** Heatmap displaying the estimated proportion of each cell type in each region by STEP. The color scale is normalized to the 0-1 range. **g** Confusion matrix comparing the transferred results with the original results (left). The spatial accuracy (sACC) by region is shown in a line graph, with the cumulative sACC displayed in a bar plot (right). **h** Spearman correlation between the in-sample and out-of-sample predictions, along with their corresponding measurement sparsity. **i** Subtype classification results at single-cell resolution for partial regions in BRCA-A and BRCA-B1. Refer to panel **j** for the legend of breast cancer subtypes. **j** Proportion of breast cancer subtypes in partially annotated regions inferred by STEP.

We next leveraged these high-resolution expression maps to dissect the molecular heterogeneity of breast cancer subtypes. By analyzing STEP-predicted expression levels of key markers including estrogen receptor 1 (*ESR1*), progesterone receptor (*PGR*), and *ERBB2*^55^, we classified the tumor regions into distinct molecular subtypes (Fig. 4i, Supplementary Fig. 61 and Supplementary Note 23). The analysis revealed inter-tumor heterogeneity: BRCA-A was dominated by Luminal B and HER2-enriched subtypes, whereas BRCA-B1 and BRCA-B2 exhibited aggressive Triple-Negative Breast Cancer (TNBC) regions alongside Luminal B cells^50,56,57^. Given that TNBC is associated with poor prognosis^58^, STEP’s ability to spatially resolve these aggressive subclones within a heterogeneous tumor mass provides critical insights.

Beyond broad subtyping, STEP’s resolution enabled a fine-grained comparison of the tumor microenvironment between invasive ductal carcinoma (IDC) and non-invasive lesions, including ductal carcinoma in situ (DCIS) and lobular carcinoma in situ (LCIS). We observed distinct spatial patterns at the tumor boundaries: the edges surrounding invasive IDC regions (tumor_edge_2) displayed a significantly more complex distribution of cancer subtypes compared to those of DCIS/LCIS (tumor_edge_1) (Fig. 4j and Supplementary Figs. 62 and 63). This suggests that the transition from non-invasive to invasive states is marked by specific alterations in the peripheral cellular neighborhood^59^. These findings highlight STEP’s capacity to not only identify cell types but also to reveal the spatial nuances of tumor invasiveness, offering potential markers for distinguishing high-risk lesions (Fig. 4i,j).

### 5. Application across diverse SRT platforms

Existing SRT techniques can be broadly categorized into two major classes: sequencing-based and imaging-based platforms. Many sequencing-based SRT platforms have relatively low spatial resolution, such as 10x Visium. In contrast, imaging-based SRT platforms, including MERFISH^60^ and osmFISH^61^, offer single-cell resolution but are restricted to a limited gene panel. Additionally, the sequencing area of many platforms is limited, such that only a subset of large tissue sections is typically profiled. In this context, STEP compensates for the limitations of diverse SRT techniques, providing a solid foundation for an array of downstream analyses (Supplementary Notes 24–26).

For sequencing-based spatially resolved transcriptomics, we evaluated STEP using a mouse cortex dataset derived from two serial sections generated by 10x Visium with spot-level resolution, whole-transcriptome coverage, and matched H&E images^29,62^. Following the workflow in Fig. 5a, we performed cell identification and gene prediction on section 2 (Fig. 5b). To further assess STEP’s predictive capability in another species under limited tissue coverage, we applied a well-trained STEP model from Section 2 to an H&E-only Section 1. Remarkably, despite using only this partial dataset, STEP demonstrated superior performance in identifying cell types and predicting gene expression, successfully revealing distinct layer-specific distribution patterns within the cortex (Fig. 5c and Supplementary Figs. 64–67).

**Fig. 5.**
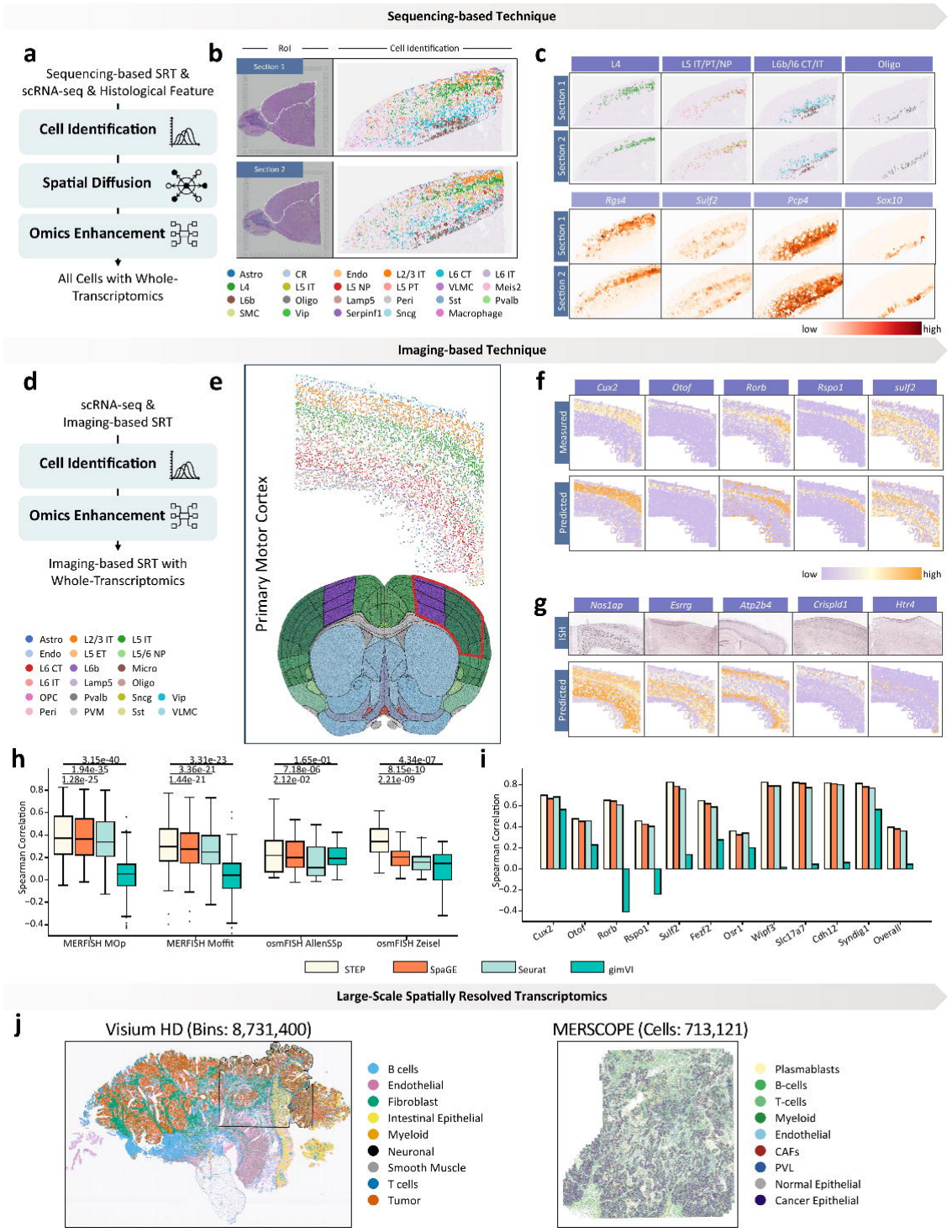
Application of STEP to diverse spatially resolved transcriptomics techniques. **a** The main workflow of inferring a spatial whole-transcriptomics at single-cell resolution from sequencing-based spatially resolved transcriptomics (SRT), single-cell RNA sequencing (scRNA-seq), and histological features through Cell Identification, Spatial Diffusion, and Omics Enhancement. **b** Hematoxylin and eosin (H&E)-stained histological image of two adjacent sections of the mouse brain cortex from the 10x Visium platform (left). White annotations indicate Regions of Interest (RoI). STEP’s cell identification results using only the gene expression data from section 2 (right). **c** Cells identification by STEP across multiple serial sections (top). The corresponding marker gene expressions, refined by STEP, are shown below (bottom). **d** The main workflow of inferring a spatial whole-transcriptomics at single-cell resolution from imaging-based SRT, and scRNA-seq through Cell Identification, and Omics Enhancement. **e** STEP’s cell identification results for the MERFISH mouse primary motor cortex dataset (top) and an anatomical reference image indicating the approximate sampling region in red (bottom). Image credit: Allen Institute. Source image: https://atlas.brain-map.org/atlas?atlas=602630314. **f** Visualization of measured and imputed expressions of layer-specific genes. **g** Visualization of imputed expression of unmeasured genes by STEP (bottom) and corresponding reference in situ hybridization (ISH) images from the Allen Mouse Brain Atlas (top). ISH images were cropped for display. Image credit: Allen Institute. Source images: *Nos1ap*, https://mouse.brain-map.org/experiment/show/77280574; *Esrrg*, https://mouse.brain-map.org/experiment/show/73616033; *Atp2b4*, https://mouse.brain-map.org/experiment/show/74735253; *Crispld1*, https://mouse.brain-map.org/experiment/show/72077456; *Htr4*, https://mouse.brain-map.org/experiment/show/72119658. **h** Boxplots displaying the Spearman correlations from the leave-one-out cross-validation experiment across four scRNA-seq datasets and their corresponding spatially resolved transcriptomics. The *p*-values above the plots are the result of performing one-sided Wilcoxon rank sum tests. The numbers of evaluated genes were *n* = 247 for MERFISH–MOp, 153 for MERFISH–Moffit, 33 for osmFISH–AllenSSp and 33 for osmFISH–Zeisel, for each method. Each observation represents one gene; genes are not independent biological replicates. For STEP versus SpaGE, Seurat and gimVI, respectively, the *p* values reported in the figure are 1.28 × □ 10 ^2^□ , 1.94 × 10□^3^□ and 3.15 × 10□ □ □ for MERFISH–MOp; 1.44 × 10□^21^, 3.36 × 10□^21^ and 3.31 × 10□^23^ for MERFISH– Moffit; 0.0212, 7.18 × 10□ □ and 0.165 for osmFISH–AllenSSp; and 2.21 × 10□ □ , 8.15 × 10□^1^□ and 4.34 × 10□ □ for osmFISH–Zeisel. Black dots represent the correlation values for individual genes. The boxplot’s center line, box limits, and whiskers denote the median, first and third quartiles, and ±1.5× interquartile range (IQR), respectively. **i** Bar plots showing the Spearman correlation of cortical layer-specific marker genes in predicting unmeasured spatial gene expression patterns. **j** Experiments on large-scale spatially resolved transcriptomics datasets, including whole-slice spatial transcriptome reconstruction on Visium HD data and genome-wide gene expression imputation on MERSCOPE data.

For imaging-based spatially resolved transcriptomics, where typically only a predefined panel of genes is measured, STEP is able to impute unmeasured genes. Here, we conducted experiments following the workflow shown in Fig. 5d on a mouse brain dataset profiled by MERFISH^63^ (Fig. 5e). Using all measured genes, STEP first inferred the cell type of each cell on the tissue slice (Fig. 5d and Supplementary Figs. 68–80). Specifically, for both 10x Visium and MERFISH data, STEP successfully reconstructed the spatial organization of the mouse cortex, showing that glutamatergic neurons exhibited strong layer specificity (*p* < 0.001 for Moran’s *I* and Geary’s *C* tests), whereas GABAergic neurons and non-neuronal cells were distributed more sparsely across layers (Supplementary Figs. 68–80).

To assess the unmeasured gene imputation performance of STEP, we performed leave-one-out imputation on genes, training on the remaining genes and imputing the spatial expression of the held-out gene. We compared STEP’s gene imputation performance with three existing methods: gimVI^20^, Seurat^21^, and SpaGE^22^ (Fig. 5h). In addition to MERFISH, we further incorporated mouse brain data of osmFISH for comprehensive evaluation^60,61,63^, with corresponding single-cell reference data including both scRNA-seq and snRNA-seq^60,64–66^. Regarding spatial expression imputation, STEP outperformed these methods by improving Spearman correlation across various SRT platforms and single-cell references (scRNA-seq and snRNA-seq) (Fig. 5h, Supplementary Table 3). Most pairwise comparisons remained significant after false discovery rate (FDR) correction (FDR < 0.05), with the only exception being the comparison between STEP and gimVI on the AllenSSp dataset (FDR = 0.165). Compared to the state-of-the-art method SpaGE, STEP showed comparable performance in imputing spatial gene expressions of cortical layer-specific markers^64^ (Fig. 5f,i and Supplementary Figs. 68 and 69). Moreover, STEP used all measured genes to impute the spatial expression of unmeasured marker genes, and the imputations were validated against in situ hybridization (ISH) images from the Allen Brain Atlas^67^, confirming their accuracy and consistency (Fig. 5g). In contrast to the analysis at the spot resolution of sequencing-based data and with a small gene panel of imaging-based data, the whole-transcriptome gene expression at single-cell resolution inferred by STEP facilitates a series of downstream analyses, providing comprehensive insights into cell-cell relationships.

We further tested the scalability of STEP on two cutting-edge SRT techniques (Fig. 5j and Supplementary Figs. 81–84): large-scale sequencing-based platforms such as Visium HD and large-scale imaging-based platforms such as MERSCOPE, which profile hundreds of thousands of cells. On the Visium HD dataset of human colorectal cancer^11^, STEP addressed the platform’s inherent constraint of limited sequencing area by reconstructing spatially resolved transcriptomes across entire tissue slices, expanding from 124,399 cells to 426,954 cells (Supplementary Figs. 81 and 82). Simultaneously, on a large-scale MERSCOPE dataset of human breast cancer containing 713,121 cells, STEP successfully imputed the genome-wide expression of unmeasured genes (Supplementary Fig. 84). It is noteworthy that, potentially due to algorithmic limitations or the lack of designed functionality for such data types, several benchmarked methods either failed to complete the analysis or could not be applied altogether.

### 6. Tri-modal integration for single-cell spatial multi-omics

To further investigate the capability of STEP in multi-omics inference for imaging-based spatially resolved transcriptomics, we performed a tri-modal integration of Xenium^52^, CITE-seq^50^, and imaging mass cytometry (IMC)^68^ data from human breast cancer (Supplementary Fig. 85). These modalities were deliberately selected for their complementary strengths: Xenium provides high-resolution single-cell information with spatial localization; CITE-seq offers paired single-cell transcriptome and surface proteome profiles, bridging the gap between RNA and protein; and IMC captures highly multiplexed protein expression. By integrating these modalities through STEP (Fig. 6a), we minimized information redundancy while enhancing cross-modality consistency. Consequently, STEP enables the construction of a single-cell spatial multi-omics map that provides a more comprehensive and biologically interpretable representation of breast cancer compared to any single modality alone.

**Fig. 6.**
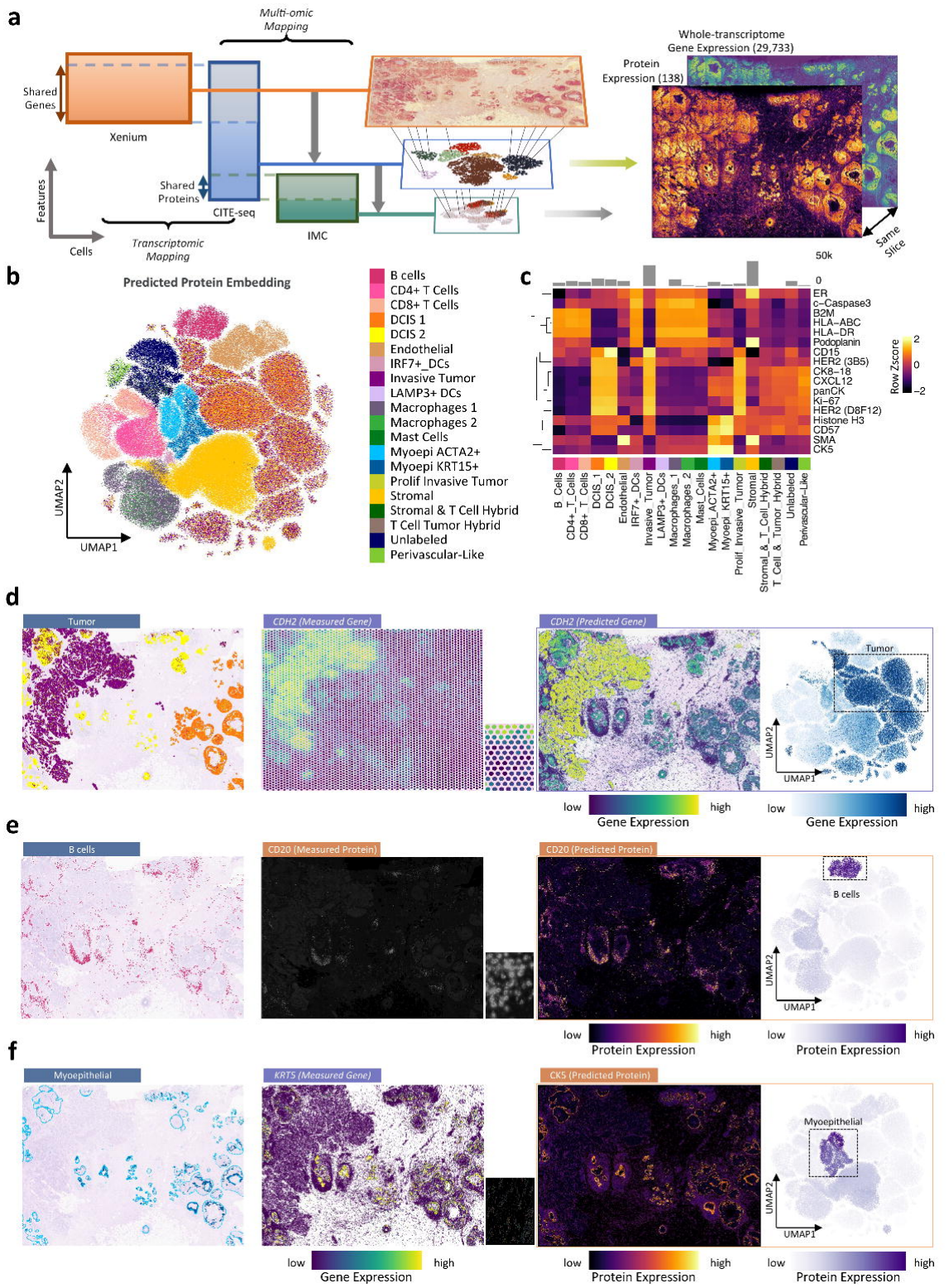
Proteomics profiling reconstruction using multi-omics reference data by STEP. **a** The main workflow of inferring spatial multi-omics atlas at single-cell resolution through Xenium, CITE-seq and imaging mass cytometry (IMC) data. **b** UMAP visualization of predicted proteomic data by STEP. **c** Heatmap displaying median proteomic profiles associated with epithelial regions across cell types. **d** Comparison among predicted transcriptomics data at single-cell resolution, the 10x Visium gene expression profile of the same name and corresponding cell types. The lower right corner of the 10x Visium gene expression profile shows an enlarged region of interest (RoI) of transcripts. The far right shows the UMAP of imputed gene. **e** Comparison among predicted proteomic data, immunofluorescence (IF) images and corresponding cell types. The lower right corner of the IF image shows an enlarged RoI. The far right shows the UMAP of predicted protein. **f** Comparison among predicted proteomic data, the gene expression profile of the same name and corresponding cell types. The lower right corner of the gene expression profile shows an enlarged RoI of transcripts. The far right shows the UMAP of predicted protein.

Using the proteomic integration result and Xenium cell-type annotations as labels, the UMAP visualization clearly showed the differences in protein expression between cancer cells and normal cells (Fig. 6b). Due to the tissue of the IMC reference not containing DCIS-type subclones, it can be observed that the model showed a mixture for cancer cell (DCIS_1, DCIS_2, invasive tumor) in UMAP (Fig. 6b), unlike normal cell types. More similar physiological conditions among the tissues of the referenced multi-omics would improve the model’s ability to distinguish different cancer subclones. We visualized the proteomic integration result for different cell types using heatmaps and found that CK5 (cytokeratin 5, a marker for basal epithelial cells) was enriched in myoepithelial cells (4.70-fold, *p* < 0.0001), whereas CD15 (a carbohydrate antigen involved in cell adhesion and signaling) was enriched in all tumor categories (2.65-fold, *p* < 0.0001) (Fig. 6c and Supplementary Fig. 86). These results were consistent with existing research conclusion^69,70^.

To more clearly characterize spatial patterns and relationships across multiple modalities, we analyzed the spatial distributions of cell types, genes, and proteins. First, STEP imputes gene expression for genes not measured in the Xenium panel, yielding expression patterns with high spatial concordance with the corresponding Visium measurements from an adjacent tissue section (Fig. 6d and Supplementary Fig. 87). We next incorporated protein-level information to further examine cross-modality relationships. CD20 is a transmembrane protein primarily expressed on the surface of mature B cells and plays a key role in regulating calcium signaling, B cell activation, and proliferation^71^. Consistent with its known expression pattern, proteomics predictions generated by STEP align with the original CD20 immunofluorescence (IF) images in the dataset (Fig. 6e). The obvious colocalization (Fig. 6c) of the inferred protein map with B cells demonstrates the robust performance of STEP. We also compared the predicted proteins with the coding gene expression of the same name, and both omics data showed consistency with their corresponding cell types (Fig. 6f and Supplementary Fig. 88). We observed variability in the concordance between predicted protein abundance and the expression of their corresponding genes: 34.88% of protein-gene pairs showed Pearson correlation coefficients greater than 0.1, whereas 37.21% showed negative Pearson correlations (Supplementary Figs. 89 and 90). This discrepancy likely stems from inherent differences between transcript-level and protein-level modalities, such as post-transcriptional regulation, post-translational modifications, and spatiotemporal shifts between mRNA production and protein localization^72,73^. Collectively, these results demonstrate that the tri-modal integration framework enables coherent integration of transcriptomic and proteomic information, supporting robust imputation of unmeasured genes and inference of protein abundance at single-cell resolution.

### 7. Quadra-modal integration for cost-effective 3D mapping

Three-dimensional (3D) spatial omics data provide advantages for deciphering biological processes by capturing spatial organization across tissue depth. However, the high cost and time-intensive nature of spatial omics experiments constrain the number of tissue sections that can be measured, especially for 3D samples. To address this limitation, we reconstructed a high-resolution 3D multi-omics brain cell atlas using just six tissue sections measured by low-resolution spatially resolved transcriptomics. This was achieved by STEP, which learns relationships between H&E image features, gene expression, and cell types from limited sections and successfully applies them to additional H&E-only sections. We collected data from 35 coronal sections across the antero-posterior axis^74^ generated by Spatial Transcriptomics, with spot-level resolution, whole-transcriptome coverage, and matched H&E images, together with snRNA-seq^25^ and inCITE-seq data^75^ for quadra-modal integration. After registering the sections to the Allen mouse brain atlas (CCFv3)^76^, we identified each cell and its corresponding gene expression within the spatial atlas using the gene expression data from only 6 sections for model training (Fig. 7a and Supplementary Note 27). Representative sections were selected to ensure that STEP was exposed to the gene expression patterns of all brain regions during training (Fig. 7b). Current methods cannot achieve super-resolution multi-omics inference to adjacent H&E-only section, making 3D multi-omics cell atlas construction particularly challenging.

**Fig. 7.**
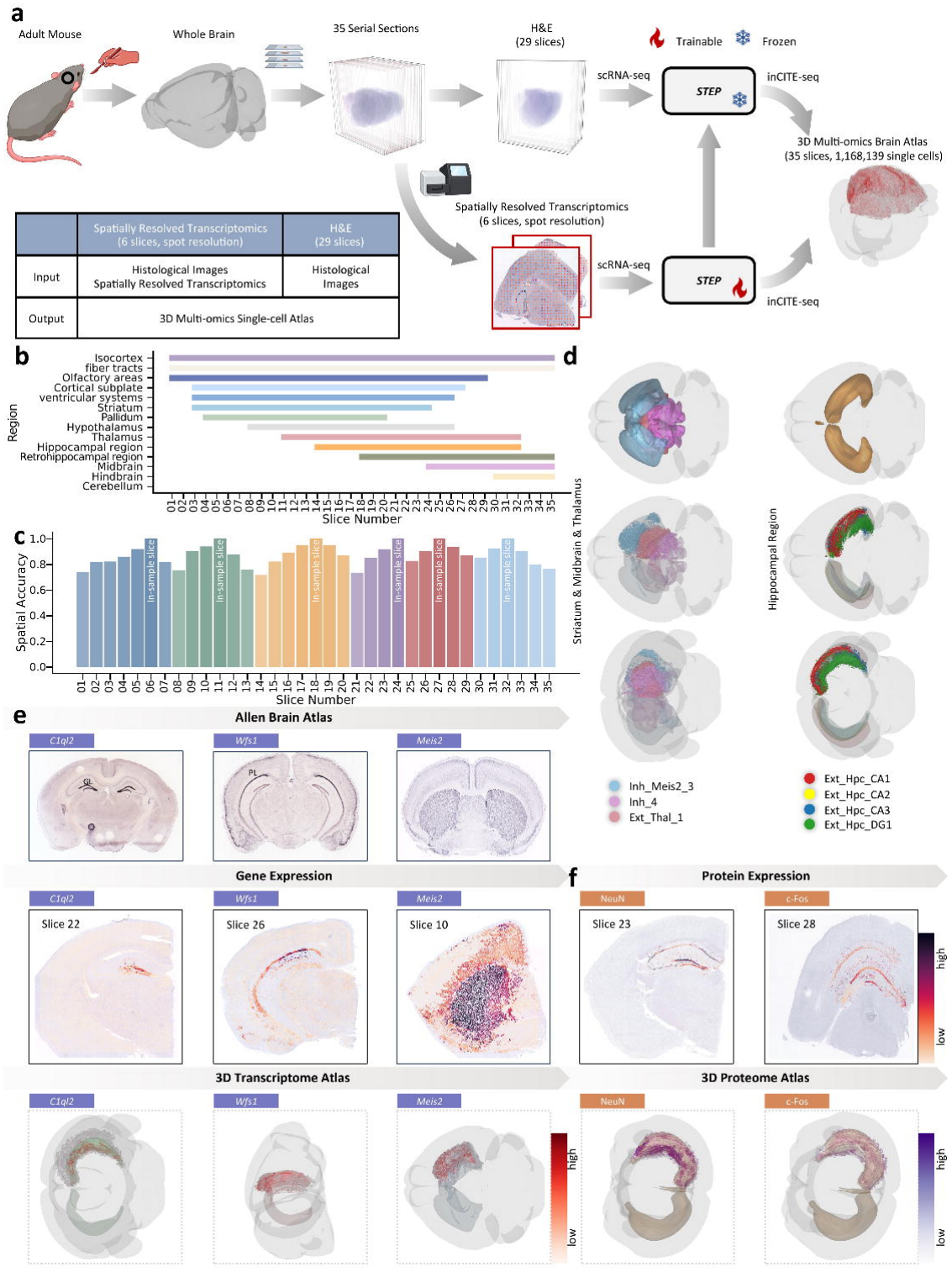
STEP’s 3D multi-omics analysis with minimum sequenced sections. **a** The main workflow of inferring a 3D multi-omics atlas with minimum sequenced sections. Sections 6, 11, 18, 24, 27, and 32 were selected as representative sections. **b** Different regions of the mouse brain in 35 coronal sections across the antero-posterior axis. **c** Bar plot showing the spatial accuracy (sACC) of 35 coronal sections between using gene expression of 6 representative sections and using gene expression of all 35 sections. Colors represent genes using the same representative sections. **d** 3D visualization of hippocampus and other subcortical structures (top). 3D visualization of the corresponding cell types identified by STEP at different angles (bottom). **e** Visualization of reference in situ hybridization (ISH) images from the Allen Mouse Brain Atlas and gene expression predicted by STEP (top). 3D visualization of predicted gene expression at single-cell resolution (bottom). ISH images were cropped and annotated for display. Image credit: Allen Institute. Source images: *C1ql2*, https://mouse.brain-map.org/experiment/show/71587929; *Wfs1*, https://mouse.brain-map.org/experiment/show/74881161; *Meis2*, https://mouse.brain-map.org/experiment/show/1231. **f** Visualization of predicted protein expression by STEP (top). 3D Visualization of predicted protein expression at single-cell resolution (bottom).

The STEP prediction, based on only 6 spatially resolved transcriptomics sections and 29 H&E-only sections, was highly consistent with the results obtained using all 35 spatially resolved transcriptomics sections. For sections without gene expression data, we observed that non-biological factors such as registration errors and staining differences increased with distance from the annotated section. Despite this, the overall spatial accuracy (sACC) remained up to 87.7%, using 6 (/35) sections with gene expression for training (Fig. 7c). Increasing the number of sections used for training STEP further improves accuracy. From the visualization results of different angles, we can see that STEP accurately identifies cells in subcortical structures such as the hippocampus and striatum (Fig. 7d and Supplementary Fig. 91).

In the mouse brain, unlike the relatively smooth surface of the cortex studied previously, the hippocampus is a curved structure with a distinct laminar organization, including the cornu ammonis (CA) regions, dentate gyrus (DG), and subiculum. Because the thickness of the pyramidal and granule layer is approximately equal to the spot diameter, the mouse hippocampus presents unique challenges for precise modeling using spatial gene expression techniques with a resolution of around 100 *μm* in diameter, particularly in relatively narrow regions like CA2^77,78^. Original Spatial Transcriptomics data blur the details of interlayer connections, whereas STEP overcomes this limitation by operating at single-cell resolution, enabling fine-grained spatial modeling of specific cortical regions. The 3D atlas of marker gene expression by STEP presents refined boundaries between different subregions of the hippocampus. Single-cell gene expression recovered the spatial patterns of various marker genes, which correctly matched the annotated hippocampal regions CA and DG in the Allen Brain Atlas^67^ (Fig. 7e and Supplementary Fig. 92). STEP’s accurate modeling of fine-grained 3D gene expression atlas builds a bridge from multi-layer coarse-grained gene expression to spatially resolved regional analysis.

Furthermore, we demonstrate the application of STEP for single-cell multi-omics studies within a 3D spatial atlas. After inferring single-cell-level gene expression in spatial contexts, STEP employs a multi-omic mapping algorithm (Methods) to construct a multi-omics map between gene and protein data (Fig. 7f and Supplementary Fig. 93). For Neuronal Nuclei (NeuN), the reconstructed protein enrichment atlas aligns well with previous studies on the distribution of mature neurons. For protein c-Fos, encoded by the immediate early gene *Fos*, the 3D reconstruction enables the investigation of the spatial and temporal dynamics of neural circuits and provides insights into the propagation of pathological activity^79^ (Fig. 7f and Supplementary Fig. 93). In summary, STEP’s attempt to reconstruct the multi-omics atlas of the mouse brain hippocampus with the help of multi-omics inCITE-seq data as a reference is successful. It shows that combining the single-omics fine-grained spatial map inferred by STEP with the reference multi-omics data to reconstruct a multi-omics spatial atlas is effective, demonstrating STEP’s excellent scalability to other omics data.

## Discussion

Spatial omics reveals cellular organization and interactions within tissues, but comprehensive multi-omics mapping at single-cell resolution remains challenging. To address this challenge, we developed STEP, a hybrid framework combining probabilistic inference and deep learning to integrate SRT, histology and single-cell reference data. It extends sequencing-based SRT to whole-section profiling at single-cell resolution and expands imaging-based SRT to whole-transcriptome coverage. Benchmarking of STEP’s basic functionality on PDAC and BRCA data demonstrated improved performance over existing methods. Through tri-modal integration of Xenium, CITE-seq and IMC, STEP mapped gene and protein expression in breast cancer at single-cell resolution. Quadra-modal integration further enabled reconstruction of a high-resolution 3D brain multi-omics atlas using gene expression data from only six sections for training, supporting cost-efficient spatial multi-omics profiling.

Although STEP demonstrates strong performance in multi-modal integration, several aspects could be further improved. Refining the distance-based assumptions of local consistency and smoothness in its omics enhancement strategy could help STEP better capture tissue heterogeneity. Developing computationally efficient generative models offers a promising direction for improving omics enhancement in large-scale applications. As spatial omics technologies advance, incorporating additional omics layers, such as epigenomics and metabolomics, could further expand STEP’s molecular coverage. Combining these measurements with computational inference could help address the resolution–throughput trade-off^2^ and advance the study of complex tissues in basic biology and precision medicine.

Gene–protein correspondence further highlights the limits of cross-modality inference. Variation in transcript–protein concordance across proteins and cell types may reflect both biological regulation and uncertainty in protein inference. Among immune-related proteins, activation markers and secreted molecules can undergo extensive post-transcriptional and post-translational regulation, whereas some lineage-defining receptors may exhibit stronger transcript–protein concordance. Transcript-based protein inference therefore captures only part of the regulatory hierarchy shaping cellular phenotypes. Incorporating additional regulatory dimensions, such as RNA velocity^80^, translational control, and proteostatic dynamics, could improve the biological realism of STEP and facilitate the study of context-dependent transcript–protein divergence across tissues.

Finally, STEP provides a framework for multi-modal integration that could support large-scale efforts such as the Human Cell Atlas and the BRAIN Initiative. By linking experimental measurements with computational inference, STEP may help reduce data acquisition requirements while revealing molecular relationships beyond individual modalities. Such integration could facilitate biological discovery and future applications in precision medicine.

## Acknowledgements

The authors have no acknowledgements to declare.

## Funding Statement

This work was supported by the National Natural Science Foundation of China (grant no. 62331011 to Yongbing Z.; grant nos. 62303119 and 32470706 to Z.Y.); the Shenzhen Medical Research Fund (grant no. D2601018 to Yongbing Z.); the Computational Biology Program of the Science and Technology Commission of Shanghai Municipality (STCSM; grant no. 25JS2850200 to Z.Y.); the Shanghai QiYuan Innovation Foundation (to Z.Y.); and the Xuemin Institute of Advanced Studies, Fudan University (to Z.Y.).

## Author Contributions Statement

Z.H., Z.Z., L.C., Z.Y., and Yongbing Z. conceptualized the study and its design. Z.H., Z.Z., L.C., Z.Y., and Yongbing Z. created the methodology. Yangen Z., X.Y., S.J., and S.D. conducted the investigation. Z.H., Yangen Z., and B.S. were involved in data collection and pre-processing. Z.H., Y.D., and J.C. performed the data analysis and visualized the study. H.W., G.W., Z.Y., and Yongbing Z. completed the project administration. Z.H. and Z.Z. drafted the manuscript. L.C., Z.W., X.L., F.W., Z.Y., Y.D., Q.Z., and Yongbing Z. critically reviewed, edited, and approved the manuscript.

## Competing Interests Statement

The authors declare no competing interests.

## Methods

### Ethics statement

This study relies on the analysis of previously published data, adhering to ethical guidelines for human and mouse samples. Data collection and analysis were not performed blind to the conditions of the experiments.

### Data pre-processing

#### Cell segmentation and feature extraction

The QuPath^81^ plugin integrated with StarDist^38,82^ was employed for cell segmentation (Supplementary Fig. 2). The StarDist model, originally introduced at MICCAI 2018, aims to train a robust nucleus segmentation framework. Pre-trained weights of the StarDist from the MoNuSeg Challenge Dataset^83^ were utilized for segmenting cells in H&E-stained images. Based on a QuPath RoI Manager tool, we expanded the nucleus masks for their cell masks, and pathologist-friendly morphological features (Supplementary Table 4) were extracted for each cell using the QuPath RoI Manager. To optimize memory usage, the large H&E images were divided into smaller tiles of 2,000 × 2,000 pixels. To prevent cells from being split at image boundaries, each tile included a 100-pixel margin overlapping with its adjacent tiles. Overlapping cells in the margins were identified based on their location distances and cell diameters. For downstream analysis, the larger cell from any pair of overlapping cells was retained.

#### Domain adaptation

In this study, we employ the Conditional Variational Autoencoder (CVAE)^40^ for generative domain adaptation to address the platform effects caused by different technologies (Supplementary Fig. 1). For example, we define *X* and *Y* as the gene expression matrices for single-cell reference data and spatially resolved transcriptomics data, adaptation to address the platform effects caused by different technologies (Supplementary Fig. 1). For example, we respectively, representing two conditions, and select *G* differentially expressed genes as informative genes. To ensure that the training data from the two conditions exhibit similar cell-type composition profiles, pseudo-spot data is generated from the single-cell reference data, providing a broad spectrum of cell-type compositions for the single-cell reference condition. The average number of cells present in a single spot, denoted as *M*, is determined based on the results of cell segmentation. Then, *M* cells are selected from the single-cell reference data and weighted according to the generated weights to compute the gene expression of a pseudo-spot. Additionally, in spatially resolved transcriptomic datasets with a limited number of real-spots, there may be significant differences in magnitude between real-spots and pseudo-spots. To address this, we randomly mix spots to generate sufficient real-spot data. Details of the CVAE training and inference procedures are provided in Supplementary Notes 2 and 28.

### Cell identification

#### Probability distribution-based inference model (Supplementary Fig. 3)

We employed a probability distribution-based inference model for cell identification. STEP integrates spatially resolved transcriptomics data and single-cell reference data as inputs. For single-cell reference data, which encompasses *T* cell types, we select genes specifically expressed in these cell types and define the *T* × *G* cell-type-specific expression matrix *θ*, where *θ*_*t,g*_ represents the mean expression level of an informative gene *g* in a specific cell type *t*. For spatially resolved transcriptomics data, the gene expression *ŷ*_*s,g*_ of gene *g* at spot *s* was inferred from *y*_*s,g*_ using CVAE. We denote *F* as the *C*×*L* morphological features matrix, where *F*_*c,l*_ represents the standardized value of feature *l* for cell *c*. The standardization is applied to each feature individually, such that for each feature, the values are adjusted to have zero mean and unit variance across all cells. These features include shape descriptors, color metrics, and other quantifiable, interpretable characteristics. We adopt an inference model based on the Poisson distribution to identify the cell type of individual cells within these spots:

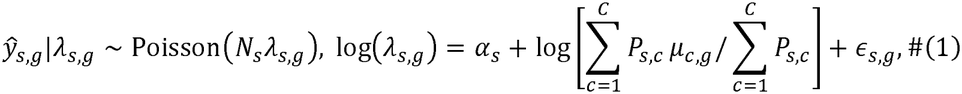

where *N*_*s*_ denotes the total number of unique molecular identifiers (UMIs) for spot *s*, and *λ*_*s,g*_ represents the relative expression level of gene *g* at spot *s*. The term *P*_*s,c*_ , derived from cell segmentation, indicates the area proportion of single cell *c* in spot *s*. We define the relative expression of an individual cell *c* for an informative gene *g* as *μ*_*c,g*_. We assume that the contribution of gene expression from single-cell *c* to spot *s* is proportional to the area proportion of this cell within the corresponding spot. Additionally, we consider the random noise 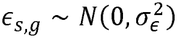, while *α*_*s*_ represents a spot-specific effect accounting for variability in a gene set across different spots.

In cell identification processes, as implemented by methods such as RCTD^84^ and SpatialScope^29^, the gene expression profiles of all cells belonging to the same type are generally assumed to be identical. However, this assumption often does not hold in real-world scenarios. Even within the same cell type, individual cells can exhibit distinct gene expression patterns that are usually associated with specific cellular characteristics. To address this limitation, we use *μ*_*c,g*_ instead of the average cell-type signature matrix *θ*_*t,g*_ :

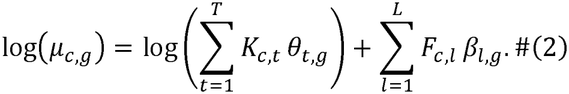

The entry *K*_*c,t*_ of the one-hot matrix *K* indicates the cell type of single cell *c*: if cell *c* belongs to cell type *t*, then *K*_*c,t*_ =1; otherwise, *K*_*c,t*_ =0. The term *β*_*l,g*,_ represents a learnable correlation matrix that captures the effect of the morphological feature *l* on the logarithm of the relative expression of gene *g* across cells. For sequencing-based SRT with matched histology images, the latter term in Equation (2) is included; otherwise, it is omitted.

In practice, the expression of a specific gene *g* is typically influenced by only a limited number of morphological features. Additionally, some pairs of morphological features may exhibit high correlation, leading to inflated regression coefficients and potentially degrading prediction performance. Therefore, we employ a spike-and-slab prior^85^ on each , *β*_*l,g*_, effectively promoting sparsity and mitigating the impact of highly correlated morphological features on regression coefficients. This approach addresses overfitting and multicollinearity while identifying significant correlations between genes and morphological features:

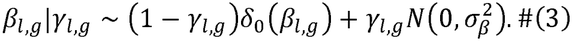

The spike component *δ*_0_ (*δ*_*l,g*_) is a point mass distribution at *δ*_*l,g*_ = 0, while the slab component 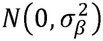 is a normal distribution with a mean of 0 and variance 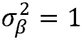. The auxiliary binary variable *γ*_*l,g*_ follows a Bernoulli distribution *Bern*(*π*_*γ*_) , where *π*_*γ*_ is governed by a Beta distribution *Beta*(*a,b*), with hyper-parameters *a*=0.5 and *b* =0.5.

When *γ*_*l,g*_ =1, *β*_*l,g*_ follows a normal distribution, indicating that the morphological feature *l* is associated with the expression of gene *g*, and the probability of *β*_*l,g*,_ being zero is effectively eliminated. Conversely, when *γ*_*l,g*_ = 0 and *β*_*l,g*_ = 0, it means that the morphological feature *l* is unrelated to gene .

#### Spatial prior

Many deconvolution methods^24,29^ have demonstrated that spatially adjacent cells are often of the same type. Moreover, in some cell segmentation tasks, cell types are identified solely based on cellular features. To leverage this information, we combine these insights into a prior Potts model^29^ that explicitly uses both morphological similarity and spatial proximity to regularize cell-type assignment. Specifically, under the assumption of spatial proximity, adjacent cells with more similar morphological features have a higher probability of sharing the same cell type. As a prior, this model incorporates the cell types and morphological features of neighboring cells to infer, for each cell, a probability distribution over cell types. For the cell type *K*_*c*_ of an individual cell *c*, we have:

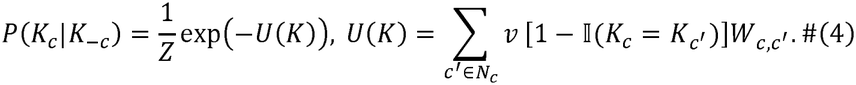

Here, *Z* is a normalization constant. The parameter *v* controls the smoothness of cell type labels, with larger values of *v* enforcing stronger spatial consistency. −*c* denotes all the cells other than cell *c. N*_*c*_ denotes the set of neighbors for cell *c*and I(·) is an indicator function that equals 1 when 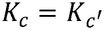 and 0 otherwise. The weight 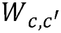 represents the similarity of morphological features between cells *c* and *c′*, quantifying their resemblance:

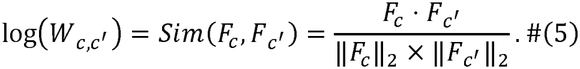

If morphology features are not available, *W* is set to a matrix of ones. Details of model fitting and parameter estimation are provided in Supplementary Note 29.

### Spatial diffusion

Spatially resolved cellular networks based on graph attention mechanisms^41^ are designed for graph-structured data. By stacking layers where nodes attend to the features of their neighbors, the networks allow for the implicit assignment of different weights to neighboring nodes. This makes spatially resolved cellular networks well-suited for semi-supervised learning tasks. In STEP, we employ a graph attention mechanism to propagate the cell types of identified cells in sequenced areas to unidentified cells in unsequenced areas in a diffusion-like manner, referred to as spatial diffusion, enabling cell identification across the whole slice and even across different slices (Supplementary Fig. 4 and Supplementary Note 30).

#### Graph Attention Network (GAT)

We first use the spatial coordinates of cells as inputs to construct a graph. In this graph, each node represents a cell, and an edge connects two cells that are physically adjacent in space. We then assign each node a feature vector derived from the cell’s morphology feature and perform semi-supervised learning, using the cell types of already identified cells as labels, to infer the cell types of all nodes, i.e., all cells in the graph. The graph structure is represented by a *C* × *C* adjacency matrix *A*, where *C* denotes the number of cells. In this matrix, 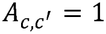if cell *c′* is among the *k*-nearest neighbors of cell *c*, and 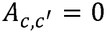otherwise. We then use the morphological feature matrix *F* and the adjacency matrix *A* to construct a GAT model *H* with multi-head attention:

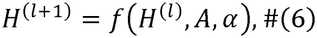

where *l* denotes the index of the graph attention layer, with *H*^(0)^ = *F* representing the initial input. The morphological feature matrix serves as the input to this model. The activations at the *l*^*th*^ layer are denoted by 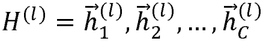 , where 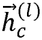 is the feature vector of cell *c* at layer *l*. The attention coefficient *α*^*(l)*^ of the *l*^*th*^ layer represents the attention weights after softmax normalization:

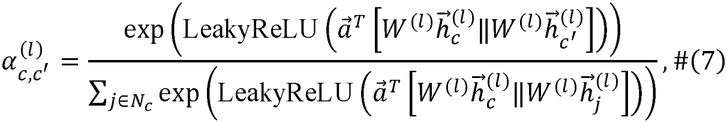

where ·*T* represents transposition and ∥ is the concatenation operation. *W*^*(l)*^ is the weight matrix of the *l*^*th*^ layer. *N*_*c*_ represents the neighbors of cell *c*. The propagation rule in the GAT can be written as:

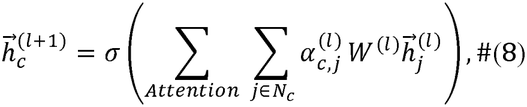

where “Attention” refers to the multi-head attention mechanism in the GAT.

The GAT model in STEP consists of multiple layers, propagating *H*^(0)^ to *H*^(4)^ , where *H*^(4)^ represents the predicted cell types for each cell, and the final layer predicts the cell types. The optimal weight matrices in the GAT can be learned by minimizing the loss of predicted labels by GAT:

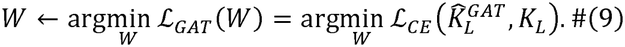

Here, ℒ_*CE*_ is a cross-entropy loss function, while *K*_*L*_ and 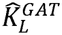 denote the inferred cell-type labels used for supervision and the corresponding GAT predictions for cells in sequenced areas, respectively.

### Omics enhancement

#### Transcriptomic mapping / Multi-omic mapping (Supplementary Fig. 5)

For sequencing-based techniques, although gene expressions were predicted by STEP at single-cell resolution, due to the gene selection process, the predicted gene expressions *μ* only contains informative genes. Prior to cell type inference, the CVAE model was used to eliminate platform effects between spatially resolved transcriptomics and single-cell reference data, and the predicted gene expressions *μ* conform to the distribution of single-cell reference data. The single-cell reference data, initially containing *U* genes before gene selection, serve as the basis for STEP to impute the expression of all spatial genes (Supplementary Note 31).

For a spatial cell *c*, we define the *k*-nearest neighbors from the *n* dissociated single-cell reference cells, using cosine distance. Next, we compute an array of *w*_*c,j*_, representing the weight between spatial cell *c* (of a certain cell type) and its nearest neighbors *j* ∈ *N*_*c*_. We retain only those neighbors with a positive cosine similarity to cell *c* (i.e., cosine distance < 1), such that:

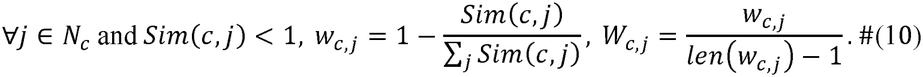

The expression 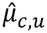 of gene *u* for cell *c* is calculated as a weighted average of the nearest dissociated cells^22^:

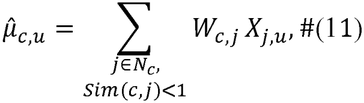

where *X*_*j,u*_ represents the expression level of gene *u* in cell *j* from the single-cell reference. For imaging-based techniques, the number of genes measured by spatially resolved transcriptomics is very limited. STEP imputes the expression of the spatially unmeasured genes from the single-cell reference.

In multi-omics settings, this strategy can be directly generalized to other pairs of omics modalities. One modality serves as the reference to define cell-wise similarity, while the other is mapped accordingly, following the same workflow described above.

#### Spatial mapping

Spatially adjacent cells exhibit similar cell types and gene expressions. Therefore, we perform spatial correction on the gene expression as obtained by spatial mapping. First, we identify *k*-nearest spots *s* ∈ *N*_*c*_ adjacent to cell *c* and construct a Gaussian kernel:

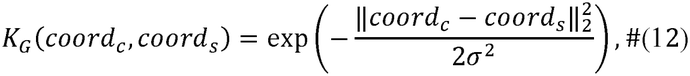

where 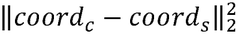 is the squared Euclidean distance, and *σ* is the length/scale parameter that characterizes the size of the focal expression patterns. The Gaussian kernel function is infinitely differentiable. Since reasonable values of *σ* yield almost identical results, we fix *σ* to be 0.1 throughout the study.

We perform spatial mapping and conduct spatial correction based on the expression of spatially resolved transcriptomics data and the Gaussian kernel function *K*_*G*_(*coord*_*c*_, *coord*_*c*_):

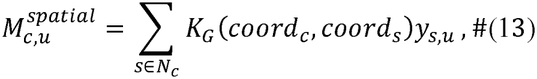

where *y*_*s,u*_ represents the expression level of gene in spot from the spatially resolved transcriptomics data. The spatial correction 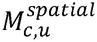 is incorporated into the gene expression data 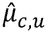 as a scaling factor:

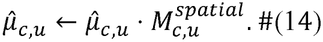

This spatial correction ensures that the gene expression data follows the inherent spatial pattern observed in the transcriptomic atlas.

### Benchmarking Comparison

Inspired by CARD^24^, for the PDAC and BRCA-A datasets, where cells lack true annotations, we use the actual region label based on H&E staining to evaluate the metrics. The true region label is assigned according to the cell’s location. The predicted labels may vary between datasets. For example, in PDAC-A, Cancer clone A and Cancer clone B cells are typically enriched in the Cancer region, so their predicted labels are updated to “Cancer”. Similarly, in BRCA-A, Cancer Epithelial cells are predominantly found in the Invasive Carcinoma region, leading to their predicted labels being adjusted to “Invasive Carcinoma”. Finally, we compute evaluation metrics such as ARI, ACC, NMI, and F1-Score using the true and predicted labels. The methods compared, their software links and parameter settings are summarized in Supplementary Table 5.

Visium-like pseudo-spots were generated by spatially aggregating single-cell Xenium profiles from human breast and colorectal cancer tissues. Using the original Xenium annotations and expression profiles as ground truth, we evaluated cell-type deconvolution with MSE and PCC, cell identification with accuracy, and gene prediction with PCC and SPCC. The colorectal cancer benchmark was also performed using an unpaired scRNA-seq reference.

### Transferring

The adaptability of GAT to dynamic graph structures enables inference of graphs with changing topologies. This characteristic facilitates the application of Spatial Diffusion, allowing a model trained on a specific tissue section to be seamlessly transferred and employed across different sections of the same tissue type. Specifically, for another tissue section, a *C′* × *C′* adjacency matrix *A′* is constructed, where *C′* represents the number of cells in the section. By applying *A′* and the corresponding morphological feature matrix *F′* to the trained model. Consequently, this approach achieves cell type identification across diverse spatial regions of the tissue.

### Implementation Details

STEP was implemented in Python (v3.10.14). Cell segmentation and morphological feature extraction were performed using QuPath (v0.5.1) with the StarDist extension (v0.5.0). The Python packages used by STEP and their versions are listed in Supplementary Table 6. Detailed model configurations, training procedures and hyperparameter settings for the conditional variational autoencoder, cell identification, spatial diffusion and omics enhancement are provided in Supplementary Notes 28–31, respectively. Experiments were conducted on a server running Ubuntu, equipped with two AMD EPYC 7763 64-core processors and 503 GiB of RAM, using a single NVIDIA GeForce RTX 4090 GPU (24 GB).

### Statistics & Reproducibility

This study is a computational reanalysis of previously published, publicly available spatial omics and single-cell datasets described in Supplementary Note 14 and the Data availability section; no new wet-lab experiments were performed. Sample sizes were determined by the availability of data in the original studies and the requirements of each analysis, with dataset sizes summarized in Supplementary Table 7. No biological samples were excluded; data preprocessing, feature selection and computational subsampling are described in the Methods and Supplementary Information. Because the study did not involve assigning samples to treatment or control groups, randomization and blinding to group allocation were not applicable. Reproducibility and robustness were assessed through repeated computational runs, leave-one-gene-out validation, transfer across tissue sections and patients, and sensitivity analyses varying reference composition, gene availability, spatial resolution, sequencing coverage and histological image style. The benchmarking analyses reported in Supplementary Tables 8–10 were repeated five times; these repetitions represent computational runs rather than independent biological replicates.

Mann–Whitney U tests were used to compare morphological feature distributions (Fig. 2d), marker protein expression across cell types (Supplementary Fig. 88) and benchmarking performance where indicated (Supplementary Tables 8–10). One-sided Wilcoxon rank-sum tests were used for comparisons of gene imputation performance (Fig. 5h) and the additional Xenium-based benchmarks (Supplementary Fig. 23). For the imaging-based gene imputation benchmarks, false discovery rate (FDR)-adjusted *p* values were used to assess significance at FDR < 0.05; gene-level *p* values were combined using Fisher’s method where indicated in Supplementary Table 3. Pearson or Spearman correlation coefficients were used to quantify agreement between molecular profiles, and Moran’s *I* and Geary’s *C* were used to assess spatial autocorrelation. Morphological differences were additionally quantified using Cohen’s and the overlap coefficient. In leave-one-gene-out validation, each target gene was withheld from the spatial input used for training and its imputed expression was evaluated against the measured spatial profile.

## Data availability

We analyzed the following publicly available datasets: (1) Human pancreatic ductal adenocarcinoma data reported in Moncada et al. (<u>GSE111672</u>); (2) Single-section 10x Visium human breast cancer data (https://www.10xgenomics.com/datasets/human-breast-cancer-ductal-carcinoma-in-situ-invasive-carcinoma-ffpe-1-standard-1-3-0); (3) Serial-section 10x Visium human breast cancer data (https://www.10xgenomics.com/datasets/human-breast-cancer-block-a-section-1-1-standard-1-1-0); (4) scRNA-seq of 26 primary tumors from three major clinical subtypes of human breast cancer reported in Swarbrick et al. (<u>GSE176078</u>); (5) Serial-section 10x Visium Mouse brain cortex data (https://www.10xgenomics.com/datasets/mouse-brain-serial-section-2-sagittal-anterior-1-standard-1-1-0); (6) Mouse brain cortex scRNA-seq reference (https://celltypes.brain-map.org/rnaseq/mouse/v1-alm); (7) MERFISH MOp dataset from the brain image library (https://doi.org/10.35077/g.8); (8) MERFISH dataset from Dryad repository (https://doi.org/10.5061/dryad.8t8s248) and the Moffit 10x dataset reported in Zhuang et al. (<u>GSE113576</u>); (9) osmFISH data reported in Linnarsson et al. (http://linnarssonlab.org/osmFISH/); (10) scRNA-seq of mouse cerebral cortex dataset reported in Zeisel et al. (<u>GSE60361</u>); (11) AllenSSp datasets reported in Wickersham et al. (https://portal.brain-map.org/atlases-and-data/rnaseq); (12) 10x Xenium human breast cancer data (https://www.10xgenomics.com/products/xenium-in-situ/preview-dataset-human-breast); (13) Human breast cancer CITE-seq reported in Swarbrick et al. (<u>GSE176078</u>) and Broad Institute single-cell portal for protein ADT information (https://singlecell.broadinstitute.org/single_cell/study/SCP1039); (14) Human breast cancer IMC data (https://zenodo.org/records/6036188#.Y2Cu8exBxqs); (15) Molecular atlas of the adult mouse brain reported in Ortiz et al. (<u>GSE147747</u>); (16) snRNA-seq of the adult mouse brain (<u>E-MTAB-11115</u>); (17) inCITE-seq reference of the mouse hippocampus reported in Chung et al. (<u>GSE163480</u>); (18) MERSCOPE human breast cancer data (https://info.vizgen.com/ffpe-showcase); (19) 10x Visium HD data and single-cell reference of colorectal cancer (https://www.10xgenomics.com/platforms/visium/product-family/dataset-human-crc); (20) Stereo-seq adult mouse brain data (https://db.cngb.org/stomics/mosta). Source data are provided with this paper.

## Code availability

The STEP software package and source code are available on GitHub (https://github.com/childishHU/STEP), together with scripts and materials for reproducing the analyses presented in this study. STEP is available as an installation-free web application at https://stepsinglecell.top/. The STEP source code is also archived on Zenodo at https://doi.org/10.5281/zenodo.23182583^86^.

## Notes

### Competing Interest Statement

The authors have declared no competing interest.

### Summary of Updates

The author list, author affiliations, and funding information have been updated.

